# Comprehensive prediction and analysis of human protein essentiality based on a pre-trained protein large language model

**DOI:** 10.1101/2024.03.26.586900

**Authors:** Boming Kang, Rui Fan, Chunmei Cui, Qinghua Cui

## Abstract

Human essential genes and their protein products are indispensable for the viability and development of the individuals. Thus, it is quite important to decipher the essential proteins and up to now numerous computational methods have been developed for the above purpose. However, the current methods failed to comprehensively measure human protein essentiality at levels of humans, human cell lines, and mice orthologues. For doing so, here we developed Protein Importance Calculator (PIC), a sequence-based deep learning model, which was built by fine-tuning a pre-trained protein language model. As a result, PIC outperformed existing methods by increasing 5.13%-12.10% AUROC for predicting essential proteins at human cell-line level. In addition, it improved an average of 9.64% AUROC on 323 human cell lines compared to the only existing cell line-specific method, DeepCellEss. Moreover, we defined Protein Essential Score (PES) to quantify protein essentiality based on PIC and confirmed its power of measuring human protein essentiality and functional divergence across the above three levels. Finally, we successfully used PES to identify prognostic biomarkers of breast cancer and at the first time to quantify the essentiality of 617462 human microproteins.

**Key Points:**

- PIC outperformed existing computational methods for predicting essential proteins.
- PIC could comprehensively predict human protein essentiality at levels of human, human cell lines and mice orthologues at the same time.
- PES could serve as a potential metric to quantify the essentiality of both human proteins and human microproteins.

## 1 Introduction

Essential genes are key components of the minimal genome required for an organism’s survival, making them indispensable **(Bartha, et al., 2018)**. Essential proteins encoded by these genes often perform core biological functions related to growth and development **(Ji, et al., 2019)**. Thus, understanding protein essentiality is crucial for identifying drug targets, clinical therapeutics, and applications in synthetic biology. Experimental methods such as single gene deletion, RNA interference, and CRISPR gene-editing technologies are commonly used to identify essential proteins **(Aromolaran, et al., 2021)**. However, these approaches are often costly, time-consuming, labor-intensive, and specific cell dependence. Therefore, there is an urgent need to develop precise and efficient computational methods for assessing protein essentiality.

Up to now, existing number of computational approaches have been developed and can be broadly classified into two main categories: network-based methods and sequence-based methods **(Aromolaran, et al., 2021)**. Network-based methods often rely on a protein-protein interaction (PPI) network and employ metrics such as betweenness centrality (BC) **(Joy, et al., 2005)**, closeness centrality (CC) **(Wuchty and Stadler, 2003)**, and degree centrality (DC) **(Hahn and Kern, 2005)** to assess the importance of each protein **(Li, et al., 2016)**. However, these methods are highly dependent on the PPI networks, which often have serious bias. Moreover, they are unable to analyze proteins that lack PPI information. In contrast, sequence-based methods are more accessible due to the central role of the sequence-structure-function paradigm. These sequence-based methods commonly utilize integrated sequence intrinsic features or biometric information to represent nucleic acid or protein sequences and then predict essentiality via machine learning models. Recently, a series of sequence-based models have been proposed, such as Pheg **(Guo, et al., 2017)**, DeeplyEssential **(Hasan and Lonardi, 2020)**, DeepHE **(Zhang, et al., 2020)**, EP-GBDT **(Zeng, et al., 2021)**, EP-EDL **(Li, et al., 2022)** and DeepCellEss **(Li, et al., 2023)**. However, the performances of these methods are limited by sequence features extraction, due to the complexity and heterogeneity of protein sequences.

More recently, the large language models (LLMs) have achieved notable success in the field of natural language processing (NLP) **(Lin, et al., 2023)**. Inspired by LLMs, protein large language models (PLMs) pre-trained on large-scale protein sequences have emerged, including ESM **(Lin, et al., 2023)** and ProtTrans **(Elnaggar, et al., 2022)**. PLMs provide a better representation of protein sequences by embedding the protein sequences in the form of high dimensional numerical vectors. With the protein representations captured by the PLMs, the performances on diverse downstream tasks, such as structure prediction **(Lin, et al., 2023)**, subcellular localization prediction **(Thumuluri, et al., 2022)**, signal peptide prediction**(Teufel, et al., 2022),** and N-linked glycosylation sites prediction **(Hou, et al., 2023)**, have been revolutionized. However, it remains unknown whether PLMs can significantly improve the tasks of essential protein prediction.

Current essential protein prediction models commonly collect data from cell viability assays for training, where essential proteins are defined as a substantial decrease in cell viability after their knockout. However, human protein essentiality is context dependent and closely related to cell type and cellular physiological stage. Yet, current methods commonly overlooked these differences and only developed general models based on integrated essentiality data rather than cell type-specific data. Besides, human essential proteins can also be identified by human genome sequencing projects and animal embryonic lethality assays like mice. At human level, proteins are defined as essential if they are rarely disturbed or truncated in humans because essential proteins are more intolerant to loss-of-function variants. While at mouse level, essential proteins are defined as human-mouse homologous proteins that cause embryonic death in mice after being knocked out. Additionally, previous studies have demonstrated that proteins identified as essential in human cell lines or knockout mice may be distinct from those in humans **(Bartha, et al., 2018)**, highlighting the urgent need for a comprehensive prediction and analysis of the human essential proteins.

Hence, for comprehensively and systematically assessing human protein essentiality across three levels (humans, human cell lines, and mice) **(Fig. 1a)**, we proposed a deep learning-based method, Protein Importance Calculator (PIC), by fine-tuning PLMs **(Fig. 1b)**, which achieved the state-of-the-art performance on human essential proteins prediction task compared to existing methods. Moreover, based on the probability values output by PIC, we defined Protein Essential Score (PES) as a potential metric to quantify the protein essentiality and measure the divergence of protein essentiality in humans, cell lines and mice **(Fig. 1c)**. Next, we confirmed the power of PES by utilizing it to identify potential prognostic biomarkers of breast cancer and quantify the essentiality of human microproteins **(Fig. 1c)**. Finally, we developed a user-friendly web server for the convenience of researchers.

**Fig. 1.**
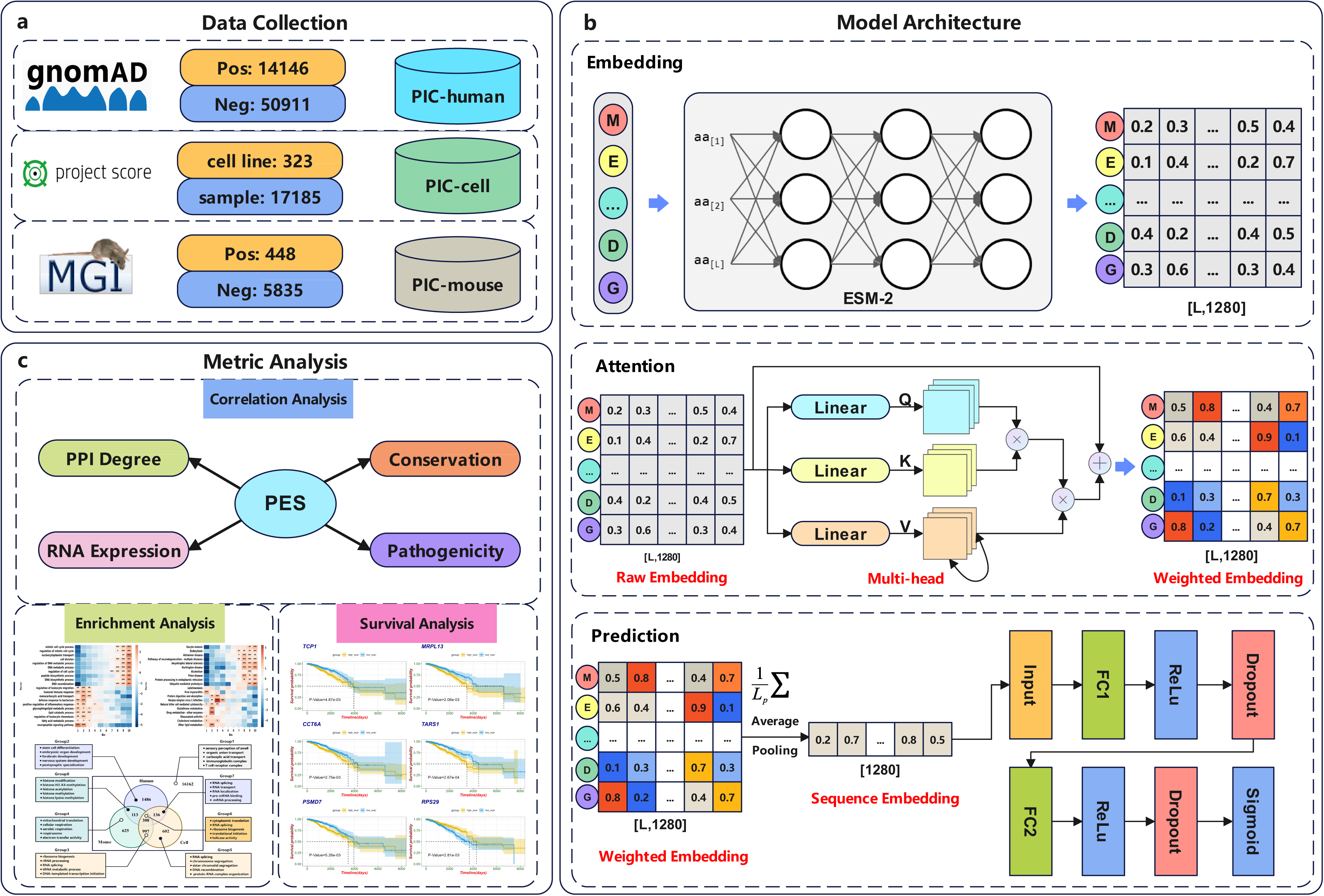
Overall workflow. (a) Data collection. Protein essentiality data were collected from gnomAD, Project Score and MGI databases to train PIC-human, PIC-cell, and PIC-mouse models, respectively. (b) Model architecture. All PIC model shared the identical architecture, including three main modules: Embedding, Attention and Prediction. The Embedding module was used to extract protein sequence features from ESM-2, the Attention module was added to capture the relative importance of amino acids at different positions in protein sequences, and the Prediction module was applied to obtain the predicted essential probability of the protein sequence. (c) Metric analysis. To confirm the power of protein essential scores (PES) generated by PIC, a series of downstream analyses of PES were conducted, including correlation analysis, enrichment analysis, and survival analysis.

## 2 Materials and Methods

In this study, a total of 325 PIC models were trained across three levels: one for humans (PIC-human), one for mice (PIC-mouse), and 323 for cell-level models (PIC-cell). Each PIC-cell model was considered as a basic classifier, and a soft voting strategy in ensemble learning was employed to aggregate the mean probability values across 323 cell lines, leading to a high-performance classifier represented by PIC-cell. Additionally, ensemble learning was also applied to construct 19 tissue-level PIC models and 28 disease-level PIC models based on the source of each cell line.

### 2.1 Data collection

As shown in **Fig. 1a**, protein essentiality data were downloaded from gnomAD, OGEE and Project Score databases to train models of PIC-human, PIC-mouse, and PIC-cell, respectively. Essential proteins were designated as positive samples, while non-essential proteins served as negative samples. More details will be illustrated in the subsequent sections.

#### 2.1.1 PIC-human

We obtained all human gene transcripts and their corresponding constraint metrics from the gnomAD database (V4.0.0) **(Chen, et al., 2024)**. Subsequently, we selected the LOEUF (loss-of-function observed / expected upper bound fraction) as the evaluation metric for protein essentiality. The LOEUF metric was calculated using data from large-scale sequencing projects conducted within the human population, representing the ratio of observed loss-of-function variants to the expected number of mutants in each gene transcript. Essential genes/proteins typically exhibit intolerance to loss-of-function variants. Consequently, the number of observed loss-of-function variants in essential gene transcripts within healthy human cohorts is expected to be significantly lower than the expected number. This implies that the LOEUF value of essential gene transcripts should be correspondingly lower **(Bartha, et al., 2018)**. Here, we employed the officially recommended LOEUF threshold as the criterion for essentiality classification. Proteins with LOEUF values below 0.6 were categorized as essential, while those with LOEUF values equal to or above 0.6 were classified as non-essential. We preprocessed the raw data obtained from gnomAD by removing missing and redundant values, and then used Ensembl BioMart **(Martin, et al., 2023)** to align the transcript IDs of genes with their respective protein sequences. As a result, we obtained 65057 protein sequences derived from 17552 protein-coding genes, along with their corresponding LOEUF values, including 14146 positive samples and 50911 negative samples.

#### 2.1.2 PIC-mouse

Given that mouse serves as a popular model organism in medicine **(Bartha, et al., 2018)**, it is quite important to investigate the essentiality of **human-mouse homologous proteins**. To achieve this, we used homologous human protein sequences as inputs for the PIC-mouse model and assigned essentiality labels based on the essentiality data of homologous mouse proteins. We retrieved mouse gene essentiality data from the Mouse Genome Informatics (MGI) database **(Eppig, 2017)**. Subsequently, we employed Ensembl BioMart to match the gene IDs with the corresponding protein sequences. This process generated 6,283 human protein sequences with their homologous mouse protein essentiality data, comprising 448 positive samples and 5,835 negative samples.

#### 2.1.3 PIC-cell

We collected the human cell line-specific binary essential score matrix from the Project Score database **(Dwane, et al., 2021)**, which were generated by Wellcome Sanger Institutes using the systematic CRISPR-Cas9 knockout screening technique in diverse human cell lines. This matrix contains varying binary essential scores for 17995 human protein-coding genes across 323 human cell lines. Then, we used Ensembl BioMart to align the gene symbols with their corresponding protein sequences. Finally, we obtained the binary essentiality labels for 17185 protein sequences across 323 human cell lines originated from 19 human tissues and 28 human diseases. Notably, the number of essential proteins varies across 323 cell lines, ranging from 353 to 2117, indicating the close relationship between protein essentiality and cell type.

### 2.2 Model architecture

PIC-human, PIC-mouse, and PIC-cell share identical model architectures, which are different only in the protein sequences and corresponding essentiality labels used as model inputs. The PIC model comprises three main modules: Embedding, Attention, and Prediction (**Fig. 1b**).

#### 2.2.1 Embedding

The purpose of this module is to extract features from protein sequences. We used a pre-trained protein large language model (PLM), ESM-2, to convert variable-length raw protein sequences into fixed-size numerical feature vectors, which is the classical embedding process. Specifically, ESM-2 has been pre-trained on over 1.8 billion protein sequences from the UniRef database using self-supervised learning, enabling it to capture intrinsic features within protein sequences **(Lin, et al., 2023)**. The parameter scale range of the ESM-2 model spans from 8 million to 15 billion. To balance the prediction performance and the computational efficiency, in this study we selected the esm2_t33_650M_UR50D model with moderate parameter size to convert the original protein sequences into numerical feature vectors.

The embedding process is summarized as follows:

A protein sequence is typically represented as:

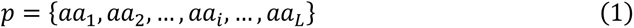

Where *p* denotes a protein sequence, *αα*_*i*_ represents amino acid residue at the *i*-th position in the sequence, and *L* is the length of the protein sequence.

Then, using the ESM-2 model, we encoded each amino acid residue in the protein sequence into a 1280-dimensional numerical feature vector, denoted as *x⃗*.

Consequently, we obtained a residue-level feature matrix *X* ∈ *R*^*L*×*d*^ to represent the entire protein sequence:

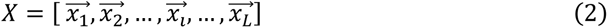

Where *L* represents the length of the protein sequence and *d* represents the dimension of embedding features.

#### 2.2.2 Attention

After the embedding module, we applied a multi-head attention module to capture the relative importance of amino acids at various positions within protein sequences. The attention module can transform the initial sequence feature matrix *X* ∈ *R*^*L*×*d*^ generated by the embedding module into a new sequence feature matrix weighted by attention weights, denoted as *X*^′^ ∈ *R*^*L*×*d*^. This new matrix incorporates information regarding the relative importance of amino acids at different positions in the sequences.

The multi-head attention mechanism is formulated as follows:

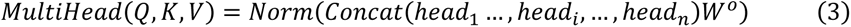

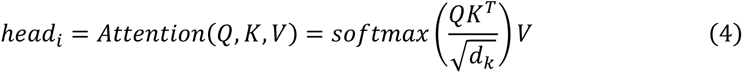

Where *head_i_* denotes the *i*-th attention head. *Q*, *K* and *V* represent query, key, and value, respectively. 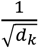 is the scaling factor of the dot-product attention. “*T*” denotes the matrix transposition. “*Norm*” refers to the layer normalization operation. “*Concat*” signifies the concatenation operation. *W*^*o*^ represents the weight matrix for output transformation.

#### 2.2.3 Prediction

In the prediction module, we employed an average-pooling layer to down-sample the high-dimensional feature representation, which converts the residue-level feature *X*^′^ ∈ *R*^*L*×*d*^ into the sequence-level feature *X*^′′^ ∈ *R*^1×*d*^ and ensures a fixed dimensionality of the model input. Then, the sequence-level feature *X*^′′^ was fed into a multi-layer perceptron to generate the raw prediction score. Finally, the raw prediction score was passed through the sigmoid activation function to obtain the predicted essential probability of the protein sequence.

### 2.3 Model details

It is well-known that essential proteins constitute approximately 10% of all human proteins **(Ji, et al., 2019)**, which will lead to highly imbalanced training datasets, which could result in prediction bias during the training process. To address this challenge, we used the WeightedRandomSampler function from the Pytorch library to enable the sampling of essential and non-essential proteins with varying weights during the training process. By adjusting these weights, we achieved a relative balanced ratio of positive and negative samples in both the training and the validation sets, thereby alleviating the learning biases during model training. Importantly, we maintained the original proportion of positive and negative samples in the test set to ensure consistency with the real situations. For all PIC models, we partitioned the dataset into 80% for training, 10% for validation, and 10% for testing. To mitigate over-fitting, we incorporated two dropout layers into the architecture: an attention dropout layer set to 0.3 and a linear dropout layer set to 0.1. In addition, we implemented a scheduler to dynamically adjust the model’s learning rate and employed an early-stopping mechanism to prevent over-fitting.

Specifically, the initial learning rate was set to 1e-5. When the loss of the validation set is no longer reduced for 2 epochs, the learning rate will be adjusted to 1/10 of the initial learning rate, and when the loss is no longer reduced for 5 epochs, the training will be terminated to prevent over-fitting.

All PIC models in this study were constructed based on Python 3.10.13 and implemented in the Pytorch 1.12.1 library. The training processes were executed on a Nvidia A100 80GB GPU. Hyper-parameter settings were determined using grid search techniques.

### 2.4 Definition of the protein essential score

Existing computational methods primarily focus on the classification of essential proteins, which overlook the exploration and analysis of model prediction probability values. The probability value generated by PIC represents the likelihood that a given protein is essential, serving as a potential quantitative metric for estimating protein essentiality. Here, we defined the probability values output by the PIC-human, PIC-mouse, and PIC-cell models as the essential scores of human proteins at the human-level, mouse-level, and cell-level, denoted as hPES, mPES, and cPES, respectively. To establish a comprehensive metric for evaluating protein essentiality, we then calculated the mean of hPES, mPES, and cPES as the protein essential score (PES) for each protein, which is used to evaluate protein essentiality across three levels. Additionally, we computed the mean of probability values output by cell-level PIC models originating from the same tissue or disease, thereby obtaining protein essential scores at the tissue-level and the disease-level.

### 2.5 Web server

We have developed a user-friendly web server that is freely accessible on the PIC website (http://www.cuilab.cn/pic). This tool allows users to input the candidate protein sequences and then obtain the essentiality of the input proteins at multiple levels. The results can be downloaded from the result page. The website was constructed base on the packages of Python 3, Flask and Numpy.

## 3 Results

### 3.1 Overall performance of PIC for essential proteins prediction

We first comprehensively predicted the human essential proteins at the human, the cell, and the mouse levels on the PIC-human, the PIC-cell, and the PIC-mouse models, respectively. Subsequently, we evaluated the performance of these PIC models on their corresponding independent test sets by Accuracy, Recall, Precision, F1 score, ROC curve area (AUROC), and PRC curve area (AUPRC) (**Table1** & **Supplementary Table S1**).

As shown in **Fig. 2a**, PIC-human achieved the highest AUROC of 0.9132, followed by PIC-mouse with an AUROC of 0.8736. The KYSE-70 cell-level model, whose AUROC (0.8579) is the median of the 323 cell-level PIC models, was selected to represent the average performance of PIC-cell models. Moreover, we categorized all PIC-cell models based on the tissue or disease source of each cell line, which further results in PIC models of 19 tissues and 28 diseases. The AUROCs of the tissue-level or disease-level PIC-cell models are ranging from 0.7543 to 0.9029 (**Fig. 2c**; **Fig. 2d; Supplementary Table S2 & Table S3**).

**Fig. 2.**
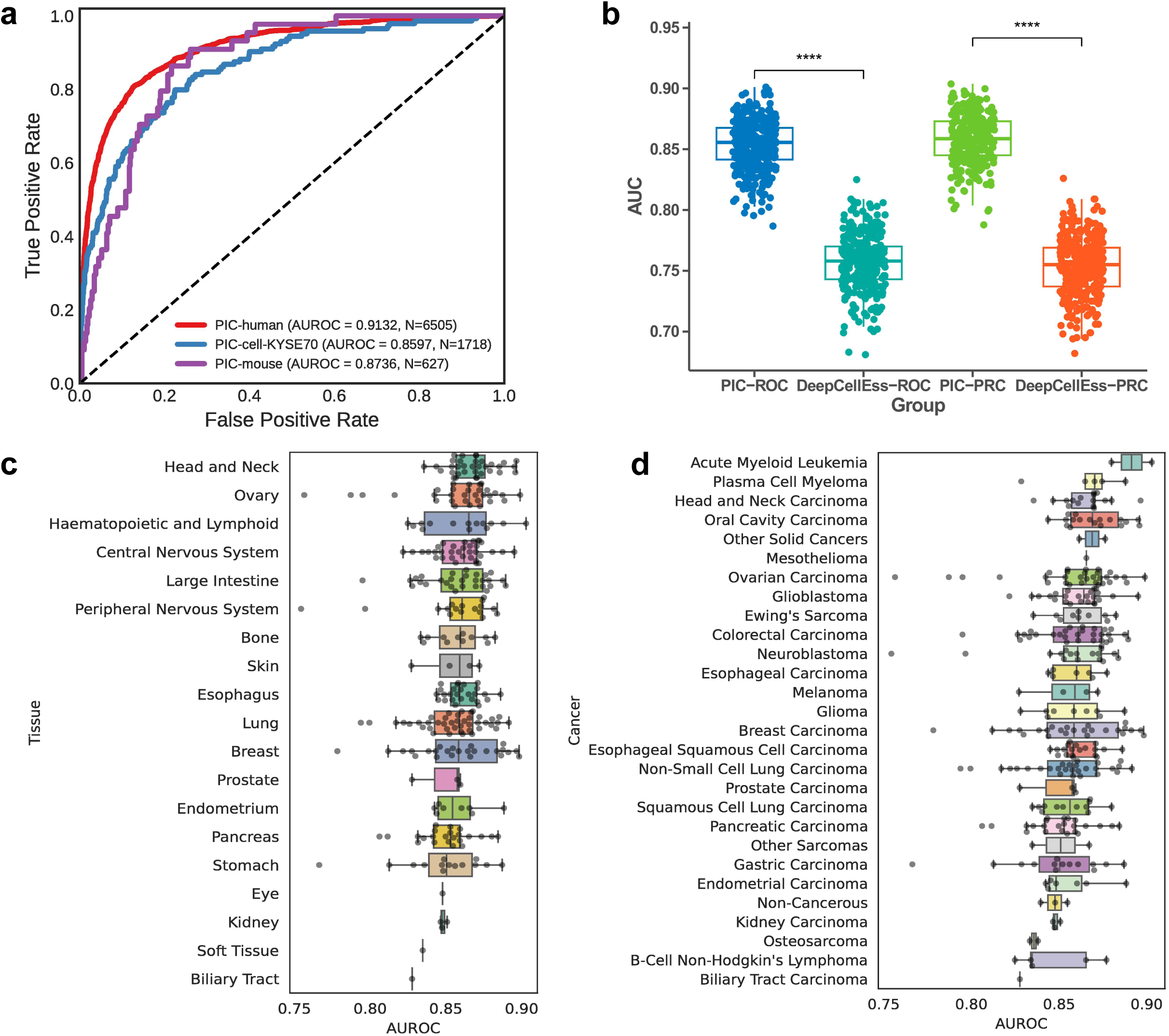
The presentation and comparison of model performance. (a) ROC curves of PIC-human, PIC-mouse, and PIC-cell models. (b) The comparison of AUROC and AUPRC between PIC and DeepCellEss on 323 human cell lines. The P-value was calculated by Mann-Whitney U rank test. *** refers to P-value<0.001. (c) The AUROCs on the independent test sets of PIC-cell models in 19 tissues. (d) The AUROCs on the independent test sets of PIC-cell models in 28 diseases.

To further evaluate the performance of PIC models, we selected three popular open-source sequence-based protein essentiality prediction models for comparison. To ensure a fair comparison of model performance, we used the training set provided by the compared methods to retrain PIC, and then compared model performance on the test set divided by the compared methods. Among the compared models, EP-EDL and EP-GBDT are both trained on an integrated dataset from cell viability assays, while DeepCellEss is a cell line-specific model based on 323 cell line datasets. Moreover, we designed PIC-base as a self-baseline model, which directly used the sequence-level feature vectors output by ESM-2 for protein essentiality prediction. To compare the performance of PIC and PIC-base, we selected protein essentiality data sourced from MDA-MB-231 cell line, which has the median number of essential proteins across 323 cell lines.

The results show that PIC increases AUROC by **5.13%-12.10%** and significantly improves Accuracy, Precision, F1 score, and AUPRC as well (**Table 2**) compared to existing methods. As DeepCellEss is a cell-line specific model, we further compared the AUROC and AUPRC between PIC and DeepCellEss on 323 cell lines one by one. As a result, compared to DeepCellEss, PIC improves the AUROC and AUPRC by an average of **9.64% and 10.52%** on 323 cell lines, respectively (**Fig. 2b**). The complete performance comparison results are provided in **Supplementary Table S4**.

**Table 1.** Overall performance of PIC models at three levels on the independent test sets.

| Model | Accuracy | Recall | Precision | F1 | AUROC | AUPRC |
| --- | --- | --- | --- | --- | --- | --- |
| PIC-human | 0.8390 | 0.8324 | 0.5924 | 0.6921 | 0.9132 | 0.7864 |
| PIC-cell | 0.8231 | 0.7135 | 0.2674 | 0.3867 | 0.8572 | 0.4277 |
| PIC-mouse | 0.8437 | 0.7045 | 0.2672 | 0.3875 | 0.8736 | 0.3064 |

Note:
- (1) The independent test sets were extreme imbalanced. - (2) The performance of the PIC-cell model was the median of 323 cell-level PIC models.

**Table 2.** Comparison of performance of PIC with existing sequence-based methods.

| Model | Accuracy | Recall | Precision | F1 | AUROC | AUPRC |
| --- | --- | --- | --- | --- | --- | --- |
| DeepCellEss | — | — | — | — | 0.7570 | 0.7532 |
| **PIC** | — | — | — | — | **0.8534** | **0.8584** |
| EP-EDL | 0.8159 | **0.8730** | 0.7474 | 0.8258 | 0.8951 | 0.8740 |
| **PIC** | **0.8587** | 0.8667 | **0.8531** | **0.8598** | **0.9375** | **0.9286** |
| EP-GBDT | 0.7300 | 0.7870 | 0.8110 | 0.7450 | 0.8150 | 0.8110 |
| **PIC** | **0.8520** | **0.8540** | **0.8510** | **0.8530** | **0.9360** | **0.9280** |
| PIC-BASE | 0.7234 | **0.6809** | 0.7442 | 0.7111 | 0.7944 | 0.7697 |
| **PIC** | **0.7570** | 0.6773 | **0.8059** | **0.7360** | **0.8457** | **0.8190** |

Note:
- (1) EP-EDL and EP-GBDT are both cell line non-specific models based, while DeepCellEss is a cell line-specific models trained on 323 human cell lines. - (2) The best performance values were highlighted in bold. - (3) In the DeepCellEss paper, only AUROC and AUPRC were calculated. The comparison between PIC and DeepCellEss was the median values of AUROC and AUPRC on 323 human cell lines.

### 3.2 PES correlates with other biological metrics of protein importance

As stated above, we defined PES output by PIC models to measure human protein essentiality. A previous study **(Chen, et al., 2020)** has reported that essential proteins are often highly expressed, conserved, disease-associated, related to core biological functions like embryo development, and inclined to act as hubs in the protein-protein interaction (PPI) network, indicating that essentiality correlates with other biological metrics characterizing protein importance. Therefore, to confirm the power of PES in measuring protein essentiality, we performed Spearman’s correlation analysis between PES and these well-established biological metrics, including PPI network node degree (STRING database), normal tissue expression level (HPA database), cancer tissue expression level (HPA database), phyloP (UCSC genome browser), phastCons (UCSC genome browser), and the number of related diseases (DisGeNet database).

As shown in **Fig. 3a**, PES has strong positive correlations with the protein degree in PPI network (Rho=0.4256, P-value<1.0e-323), normal tissue expression level (Rho=0.3495, P-value<1.0e-323), cancer tissue expression level (Rho=0.4945, P-value<1.0e-323), phyloP (Rho=0.4585, P-value<1.0e-323), phastCons (Rho=0.4081, P-value<1.0e-323), and the number of related diseases (Rho=0.1912, P-value=9.3e-139). In addition, we ranked human proteins by PES in ascending order and then divided them into 10 equal-sized groups to observe the trend of various biological metrics from group1 to group10. The results also showed robust positive correlations between PES and all of the observed biological metrics (**Fig. 3b & Supplementary Figure S1**).

**Fig. 3.**
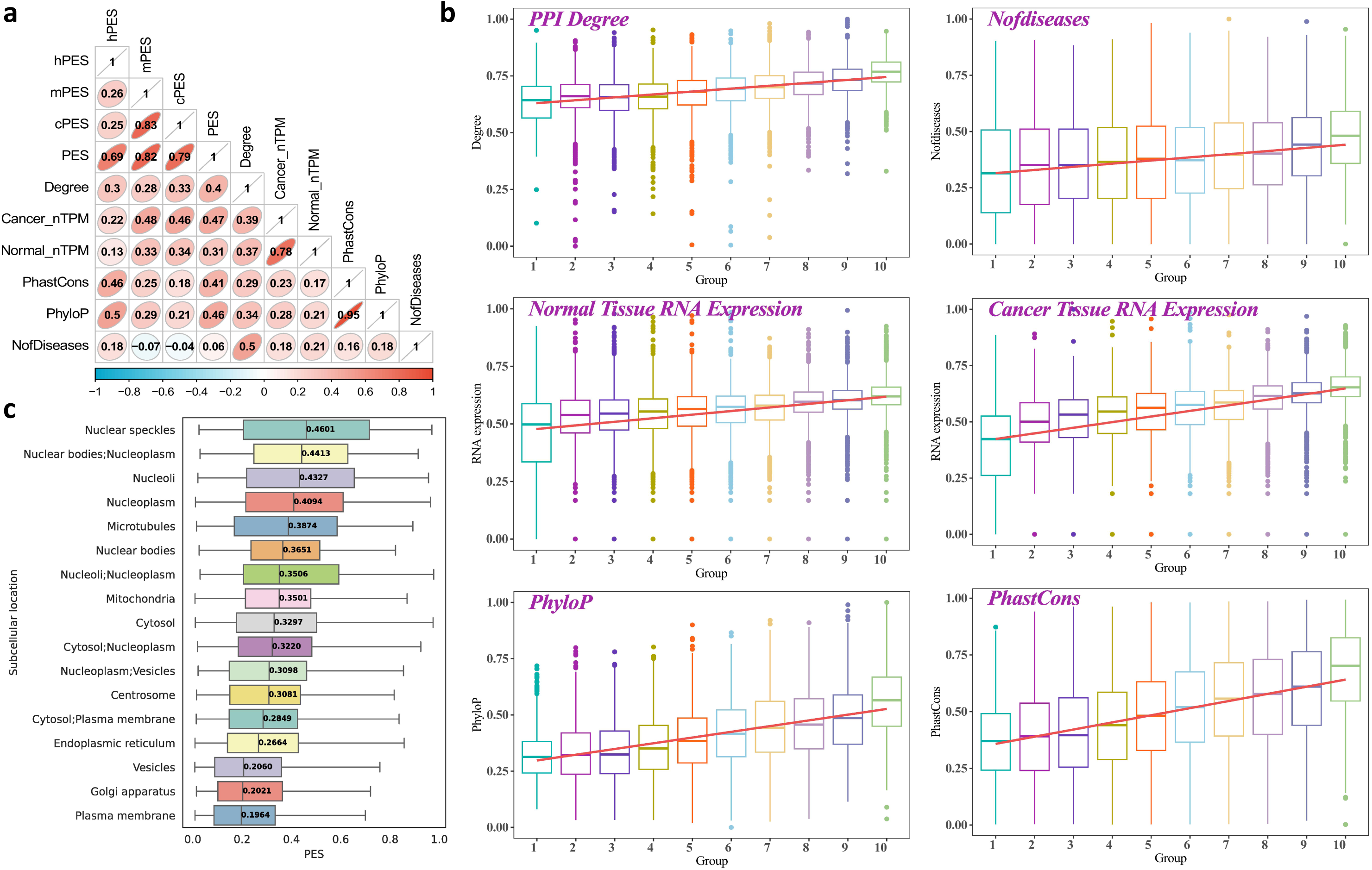
The Spearman’s correlation analysis between PES and other well-known biological metrics characterizing protein importance. (a) The spearman’s correlation coefficient matrix between PES and other biological metrics. The color intensity and numerical values in the figure reflect the strength of the correlation between any two metrics. (b) The detailed correlation analysis results between PES and 6 well-known protein importance metrics. (c) The distribution of PES across proteins with distinct subcellular localizations.

It is well-known that essential proteins tend to perform crucial biological functions at corresponding subcellular localization. We further grouped human proteins based on their main subcellular localization categories based on HPA database annotation and then calculated PES for proteins in each category. As shown in **Fig. 3c**, proteins with higher essentiality tend to localize closer to the cell nucleus, while those closer to the cell membrane tend to have lower PES. Further, we divided proteins into two groups based on whether they are localized at the cell nucleus. Then, we conducted a chi-square test for all human proteins between the binary essentiality defined by PES and the binary subcellular localization. The result also shows that essential proteins tend to locate at the cell nucleus, while non-essential proteins tend to locate outside the cell nucleus (Chi-square test; P-value=3.42e-66). These observations are consistent with the fact that vital cellular processes like replication, transcription, and translation prefer to occur near the cell nucleus.

The above results suggest strong correlations between PES and other metrics representing protein importance across various dimensions, highlighting PES as a potential metric for measuring protein essentiality.

### 3.3 Identifying essential functions using PES

Given that PES serves as a powerful metric for assessing protein essentiality, it is thus feasible to identify essential functions performed by essential proteins using PES. For this purpose, we investigated the relation between PES and protein functions. We first sorted human proteins by PES in ascending order and then divided them into ten equal-sized bins. Subsequently, we performed functional enrichment analyses on proteins within each bin, including Biological Process (BP), Cellular Component (CC), Molecular Function (MF) and Kyoto Encyclopedia of Genes and Genomes (KEGG). Then, we aggregated annotation terms from the ten bins, focusing on terms shared by all bins to observe relations between functions and essentiality. Odds ratio (OR) was calculated for each annotation term and then transformed via log_2_, denoted as log_2_OR. Spearman’s correlation coefficients were then calculated between each term’s log_2_OR and the bin number across ten bins. We selected the top 10 terms with the highest positive and the negative correlation coefficients to show the relation between functions and essentiality.

As shown in **Fig. 4a**, proteins with higher PES tend to be more associated with essential biological processes like cell division, DNA biosynthetic process, and peptide biosynthetic process. In contrast, proteins involved in more complex functions such as humoral immune response, neuropeptide signaling pathway, and fatty acid metabolic process tend to have lower PES. Moreover, CC and MF enrichment results are available in **Supplementary Figure S2 & Figure S3.** As for KEGG, we observed that highly essential proteins tend to be linked to neurodegenerative diseases such as Alzheimer’s disease, Huntington’s disease, and prion disease, while highly non-essential proteins tend to be connected to functions related to digestion and metabolism, including Protein digestion and absorption, Drug metabolism, and Cholesterol metabolism (**Fig. 4b**). The detailed enrichment results are available in **Supplementary Table S5 & Table S6.**

**Fig. 4.**
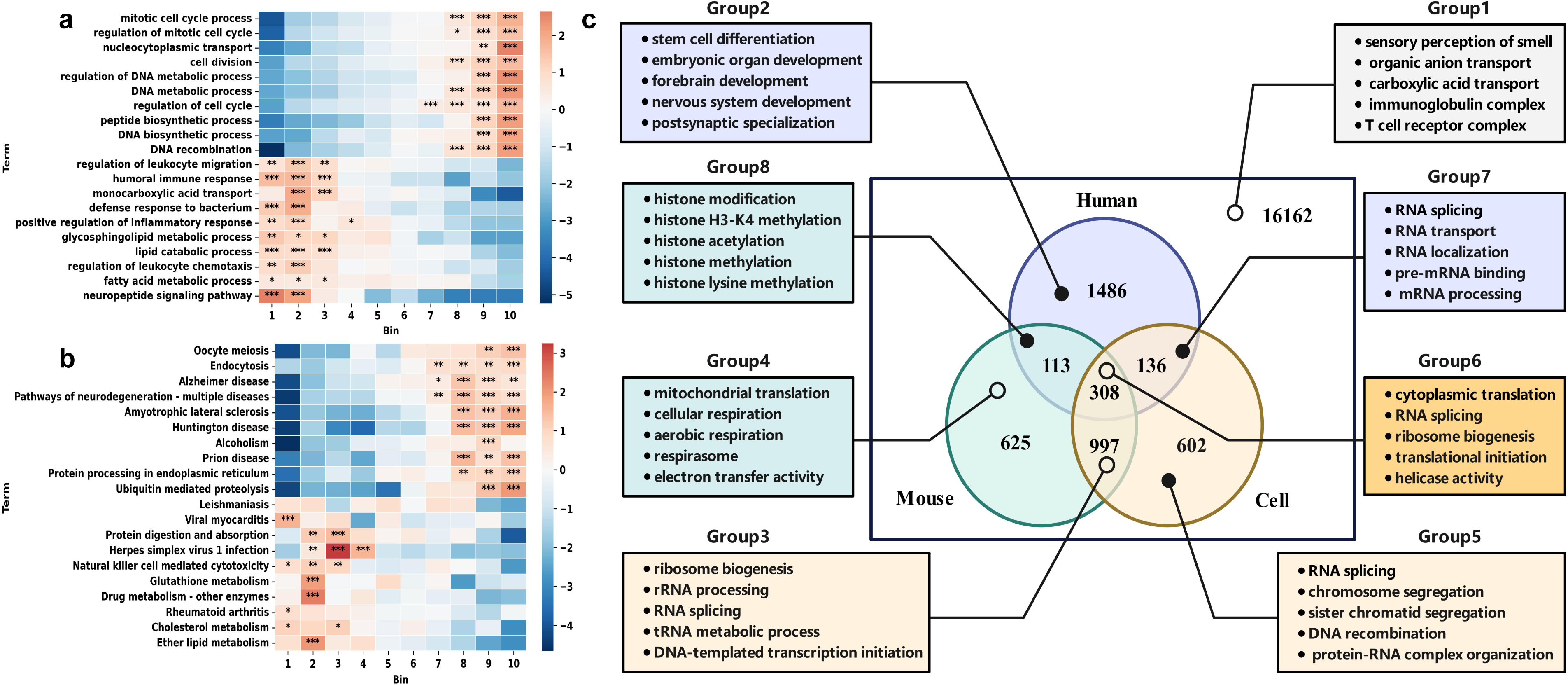
Functional enrichment analysis results of proteins with different PES and differentially essential proteins (DEPs). (a) Biological process (BP) enrichment analysis results of 10 protein bins with different PES. (b) KEGG enrichment analysis of 10 protein bins with different PES. (c) Functional enrichment analysis results of DEPs in 8 different groups. The color scale represents the log_2_OR value of each term. * refers to p<0.05; ** refers to p<0.01; *** refers to p<0.001. The P-value was calculated by Fisher’s exact test.

The overall results suggest that PES is highly correlated with protein functions. As it increases, protein functions are more associated with essential biological processes like DNA replication, protein synthesis, and cell division. Conversely, as the PES decreases, protein functions tend to be more involved in complex biological roles such as immune responses, nutrient metabolism, and nervous system maintenance. This highlights the potential of PES as a potential metric for measuring protein function essentiality.

### 3.4 Discovering functional divergence of essential proteins across three levels via PES

Due to the variation of protein essentiality across humans, human cell lines, and mice, we defined hPES, cPES, and mPES to measure protein essentiality at human-level, cell-level, and mouse-level, respectively. As shown in **Fig. 3a**, there is a strong correlation between mPES and cPES (Rho=0.83, P-value<1.0e-323), while hPES has weaker correlations with mPES (Rho=0.26, P-value=5.0e-291) and cPES (Rho=0.25, P-value=5.5e-290). Furthermore, we performed Spearman’s correlation analysis between 323 cell-level cPESs and hPES, and then sorted all cell lines by their correlation coefficients with hPES. The results showed that the ovarian cancer cell line OAW-42 had the highest correlation with hPES (Rho=0.3971, P-value<1.0e-323), whereas the large cell lung carcinoma cell line LCLC-97TM1 had the lowest correlation (Rho=0.1146, P-value=5.2e-59, **Supplementary Table S7**). These results are consistent with the divergence of protein essentiality across three distinct levels, indicating the necessity of assessing protein essentiality comprehensively via different levels of PES. Furthermore, we tried to discover the functional divergence of essential proteins across three levels via PES. Firstly, we defined Differentially Essential Proteins (DEPs) as the proteins that have different labels predicted by PIC at the human, mouse and cell-level. Proteins ranked in the top 10% sorted by hPES, mPES, and cPES are considered essential at each level, while the rest proteins are considered non-essential. Therefore, 8 groups of DEPs were generated by combining predicted labels from the three levels (2*2*2=8). Subsequently, GO and KEGG enrichment analyses were performed on the DEPs in the 8 groups for functional annotation, and the representative terms were shown in **Fig. 4c**. The specific enrichment analysis results are available in the **Supplementary Table S8 & S9**. The results showed that DEPs in different groups perform diverse functions (**Fig. 4c**). For example, by comparing the enrichment analysis results between group 2 and group 3, it is apparent that proteins essential only in humans (group 2) play a key role in maintaining organogenesis and development in complex systems, especially in the nervous system. In contrast, proteins essential in both mouse and human cell lines (group 3) are involved in genetic information transmission and providing energy for cell proliferation.

Given that most of the cells in humans are highly differentiated with low proliferative capacity while those in mouse embryos and human cell lines have high proliferative activity, it is no wonder that the essential proteins identified at different levels carry out diverse functions. Therefore, it further suggested that, for humans, human cell lines, and mice, the difference in cellular physiological stage results in the divergence in functional requirements, and then leads to the divergence in protein essentiality. Hence, PESs derived from different levels of PIC could serve as a powerful metric to identify the functional divergences.

### 3.5 Case studies

#### 3.5.1 Finding potential prognosis biomarker of breast cancer through PES

Breast cancer is the most common malignancy among women, posing a formidable health challenge worldwide **(Jabbarzadeh Kaboli, et al., 2024)**. And it is thus important to develop biomarkers efficiently predicting the prognosis of breast cancer. To test the potential of the proposed PES algorithm in this task, we collected the TCGA-BRCA project data from the cancer genome atlas (TCGA) database. We downloaded patients’ transcriptome data and clinical information from TCGA. After filtering out genes with low-level expression and patient data with incomplete clinical survival information, we ultimately obtained the expression profile data of 19,938 genes for 1,115 breast cancer patients. Employing an ensemble learning strategy, we integrated the 25 PIC models trained on breast cancer cell lines into a general breast cancer level PIC model, denoted as PIC-BRCA. Furthermore, by using PIC-BRCA, we calculated the PESs in breast cancer, denoted as PES-BRCA. Proteins with higher PES-BRCA are expected to be more essential for the survival of breast cancer cells, which then could serve as potential therapeutic targets or prognostic biomarkers. We ranked all proteins by PES-BRCA, and then selected the top 10 proteins with highest PES. Subsequently, we categorized the 1,115 breast cancer patients into high-expr and low-expr groups based on the median expression level of the 10 proteins in the patients. Log-rank tests were performed between the high-expr and the low-expr groups to investigate whether the 10 proteins are associated with the survival of breast cancer. As shown in **Table 3 & Fig. 5**, all of the 10 proteins are able to predict the survival of breast cancer (Log-rank test, P-value<0.01). In addition, 6 out of the 10 proteins have been supported by publications. This case study indicates that PES is able to discover prognostic biomarkers or even therapeutic targets. It is thus expected that PES could assist in finding prognostic biomarkers and therapeutic targets in other types of cancers and disease because we can combine various cell-level PIC models to measure protein essentiality as needed.

**Fig. 5.**
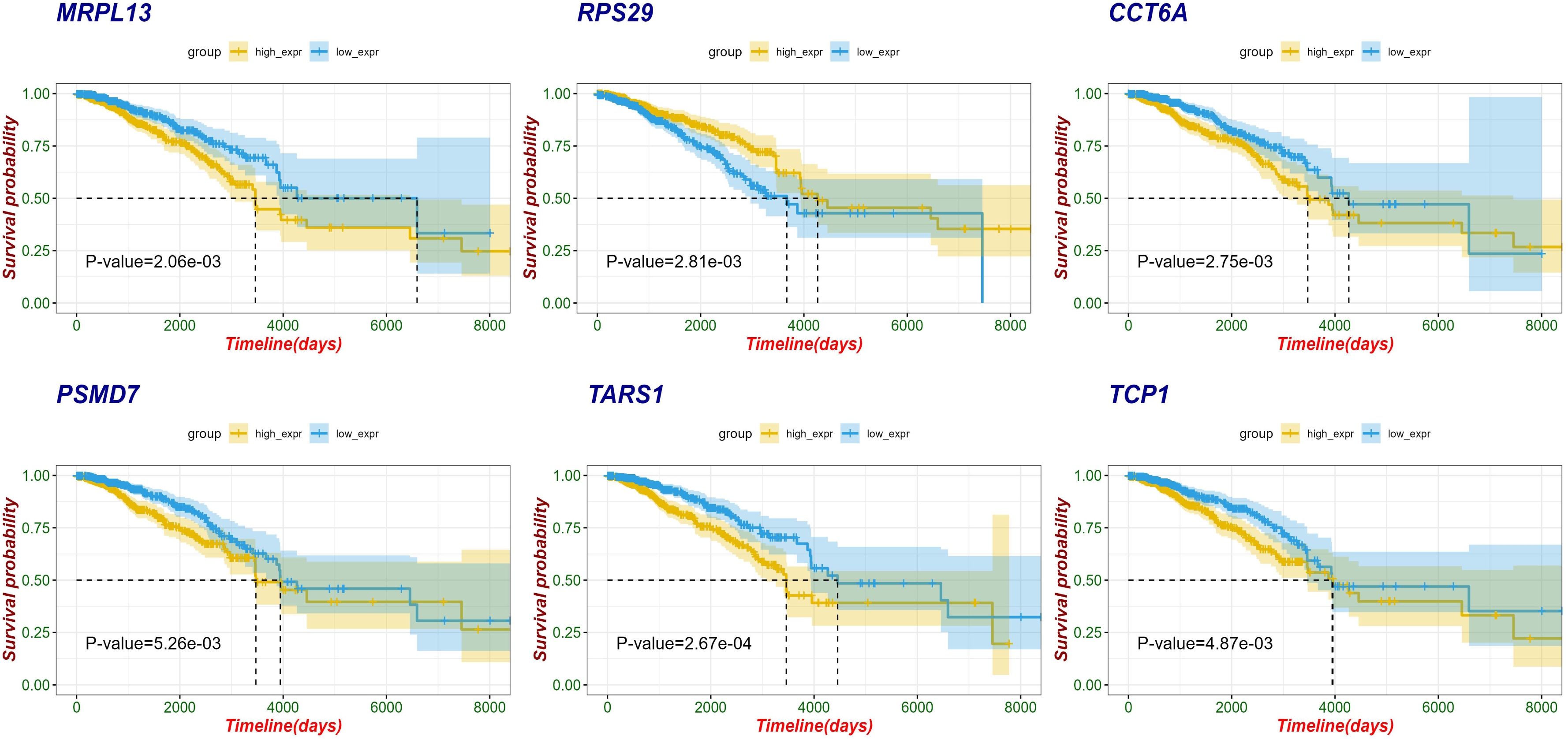
The potential prognosis biomarkers for breast cancer selected via PES-BRCA. The 1115 breast cancer patients were categorized into high-expr (yellow curve) groups and low-expr (blue curve) groups based on the median expression level of these selected proteins. The P-value was calculated by the log-rank test.

**Table 3.** 10 potential prognosis biomarkers for breast cancer screened by PES-BRCA.

| Protein | PES-BRCA | P-value | PMID |
| --- | --- | --- | --- |
| RPF2 | 0.9814 | 0.0049 | NA |
| RPS24 | 0.9724 | 0.006 | NA |
| SDAD1 | 0.9639 | 0.0025 | NA |
| TCP1 | 0.9636 | 0.0049 | 37539774 |
| MRPL13 | 0.9529 | 0.0021 | 33658859 |
| GARS1 | 0.9480 | 0.00072 | NA |
| PSMD7 | 0.9454 | 0.0053 | 33042429 |
| CCT6A | 0.9427 | 0.0027 | 37190091 |
| TARS1 | 0.9355 | 0.00027 | 37637061 |
| RPS29 | 0.9265 | 0.00281 | 37198999 |

Note: P-value was calculated by the log-rank test

#### 3.5.2 Quantifying human microprotein essentiality via PES

In the human genome, there are approximately 20,000 protein-coding RNAs, but there are millions of microproteins translated from small open reading frames (smORFs), which are open reading frames (ORFs) with lengths within 100 codons **(Ji, et al., 2020)**. Existing literature has confirmed that microproteins play crucial roles in regulating cell proliferation **(Polycarpou-Schwarz, et al., 2018)**, cellular respiration **(Makarewich, et al., 2018)**, and immune modulation **(Bhatta, et al., 2020)**. However, there are still no methods available for measuring the essentiality of microproteins up to now. It is thus important to test whether PES is able to evaluate human microprotein essentiality or not.

**Ji et al.** proposed smORFunction, a tool that integrates 617,462 unique smORFs and their corresponding microproteins, for the functional prediction of these microproteins **(Ji, et al., 2020)**. We obtained the sequences of all human microproteins from smORFunction and employed the PIC models to predict essentiality of the 617,462 microproteins by calculating their corresponding PESs. To confirm the efficacy of PES in measuring microprotein essentiality, we selected the top 10 microproteins with highest PESs **(Table 4)** and then used smORFunction to predict the functions of the 10 microproteins. The results showed that these microproteins mainly involves in essential biological processes such as cell division, cellular respiration, and DNA replication **(Supplementary Table S10)**, which is consistent with that for essential proteins. This results further suggest that PES could be also used to efficiently quantify the essentiality of microproteins.

**Table 4.** Top 10 microproteins with the highest PES predicted by PIC models.

| Microprotein | Length | hPES | mPES | cPES | PES |
| --- | --- | --- | --- | --- | --- |
| smORF594570 | 77 | 0.9925 | 0.9477 | 0.9836 | 0.9746 |
| smORF594569 | 59 | 0.9868 | 0.9243 | 0.9608 | 0.9573 |
| smORF597959 | 100 | 0.9870 | 0.9133 | 0.9234 | 0.9413 |
| smORF597276 | 99 | 0.9137 | 0.9320 | 0.9666 | 0.9374 |
| smORF614125 | 94 | 0.8883 | 0.9295 | 0.9791 | 0.9323 |
| smORF614698 | 97 | 0.8920 | 0.9253 | 0.9545 | 0.9239 |
| smORF605028 | 93 | 0.9245 | 0.8776 | 0.9566 | 0.9196 |
| smORF598483 | 79 | 0.9846 | 0.8173 | 0.9458 | 0.9159 |
| smORF590947 | 52 | 0.9467 | 0.8483 | 0.9437 | 0.9129 |
| smORF587640 | 81 | 0.9788 | 0.7761 | 0.9535 | 0.9028 |

## 4 Discussion and conclusion

No protein is absolutely essential; only functions can be so **(Chen, et al., 2020)**. Human protein essentiality is context dependent and closely associated with cell type and physiological stage. However, existing computational methods inadequately consider the divergence of protein essentiality across humans, human cell lines, and mice, just predicting protein essentiality at human cell lines level. Therefore, we proposed a deep learning framework, PIC, by fine-tuning a protein large language model and defined PES from three levels to measure human protein essentiality comprehensively. As a result, we confirmed that PIC significantly outperformed existing computational methods.

PES is the probability value output by PIC and serves as a potential metric for evaluating protein essentiality. Furthermore, we explored the power of PES from multiple aspects. First, PES is highly correlated with other well-known protein importance metrics, indicating that PES could characterize protein importance. Moreover, PES is useful for identifying essential functions. It is shown that proteins with higher PES tend to involve in core biological functions like cell division, cellular respiration, and DNA replication. In addition, PES is able to decipher the functional divergence of essential proteins identified in humans, human cell lines, and mice. Moreover, we showed that PES is also able to find potential prognostic biomarkers for breast cancer and potentially for other cancers and diseases. Additionally, PIC represents the first tool for quantifying the essentiality of human microproteins. Finally, we developed a user-friendly web server for the convenience of researchers. We believe that PIC will be beneficial for users to comprehensively predict and understand the essentiality of human proteins and microproteins, which is helpful for the discovery of therapeutic targets and prognostic biomarkers.

There also exist some limitations in the current study. For example, we defined PES using the probability values outputted by PIC and conducted preliminary exploration and analysis of its biological meaning. However, we lack in-depth explanation of the core biological meaning of PES, which is largely due to the fact that the neural network model is a black box. Fortunately, with the rapid development of interpretability in deep learning models, we believe that in the future, we can further explore the biological meaning of PES and apply them to solve broader biomedical problems.

## ACKNOWLEDGEMENTS

This study was supported by the grants from the National Natural Science Foundation of China [62025102, 81921001] and the Scientific and Technological Research Project of Xinjiang Production and Construction Corps (2023AB057).

## Author contributions

QC presented the original idea. BK and RF designed the study. BK performed the study. BK, RF, CC and QC wrote or edited the manuscript.

