## Supplementary figures and images for "Comprehensive prediction and analysis of human protein essentiality based on a pre-trained protein large language model"

### Supplemental Figure 1

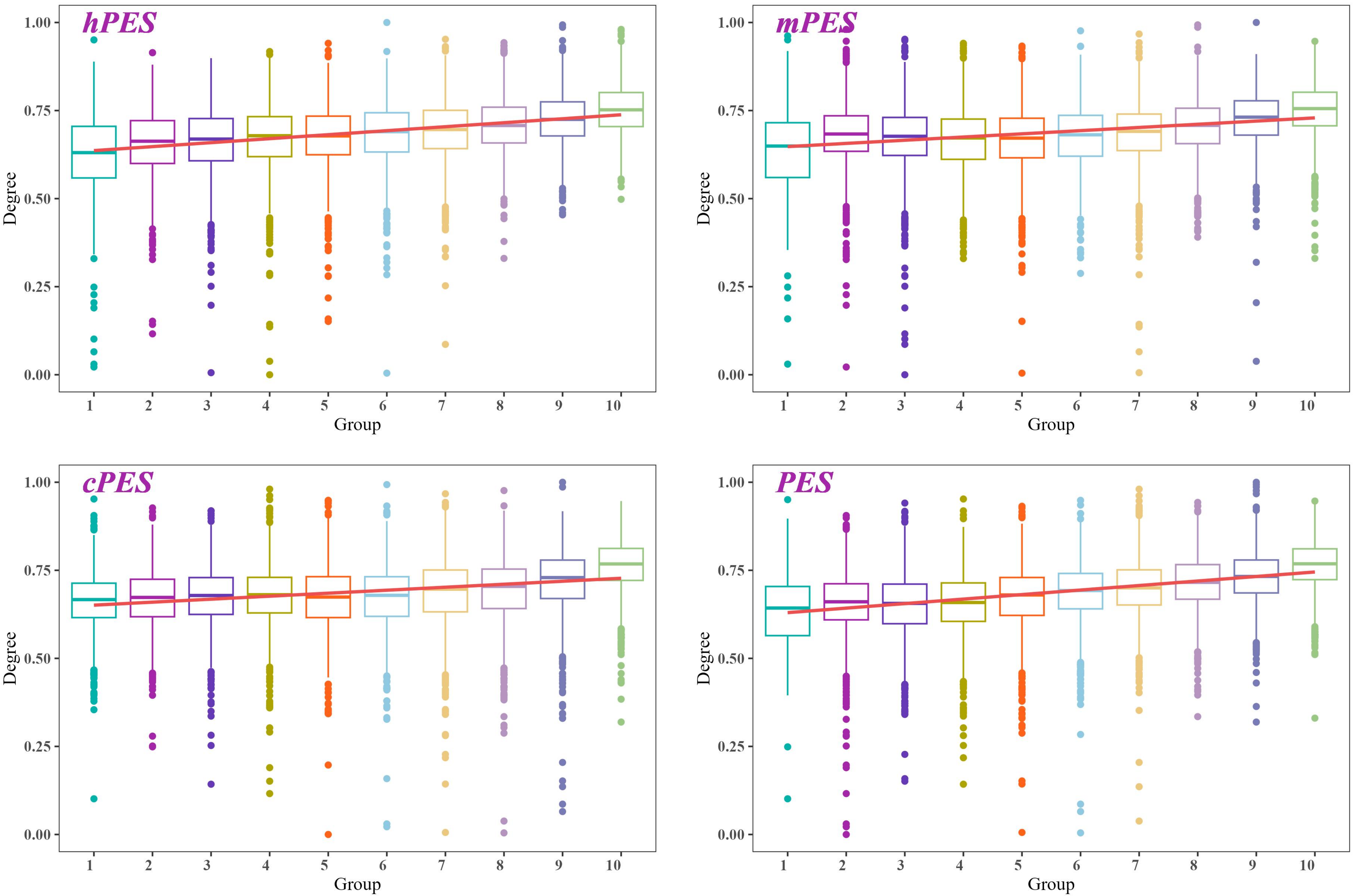

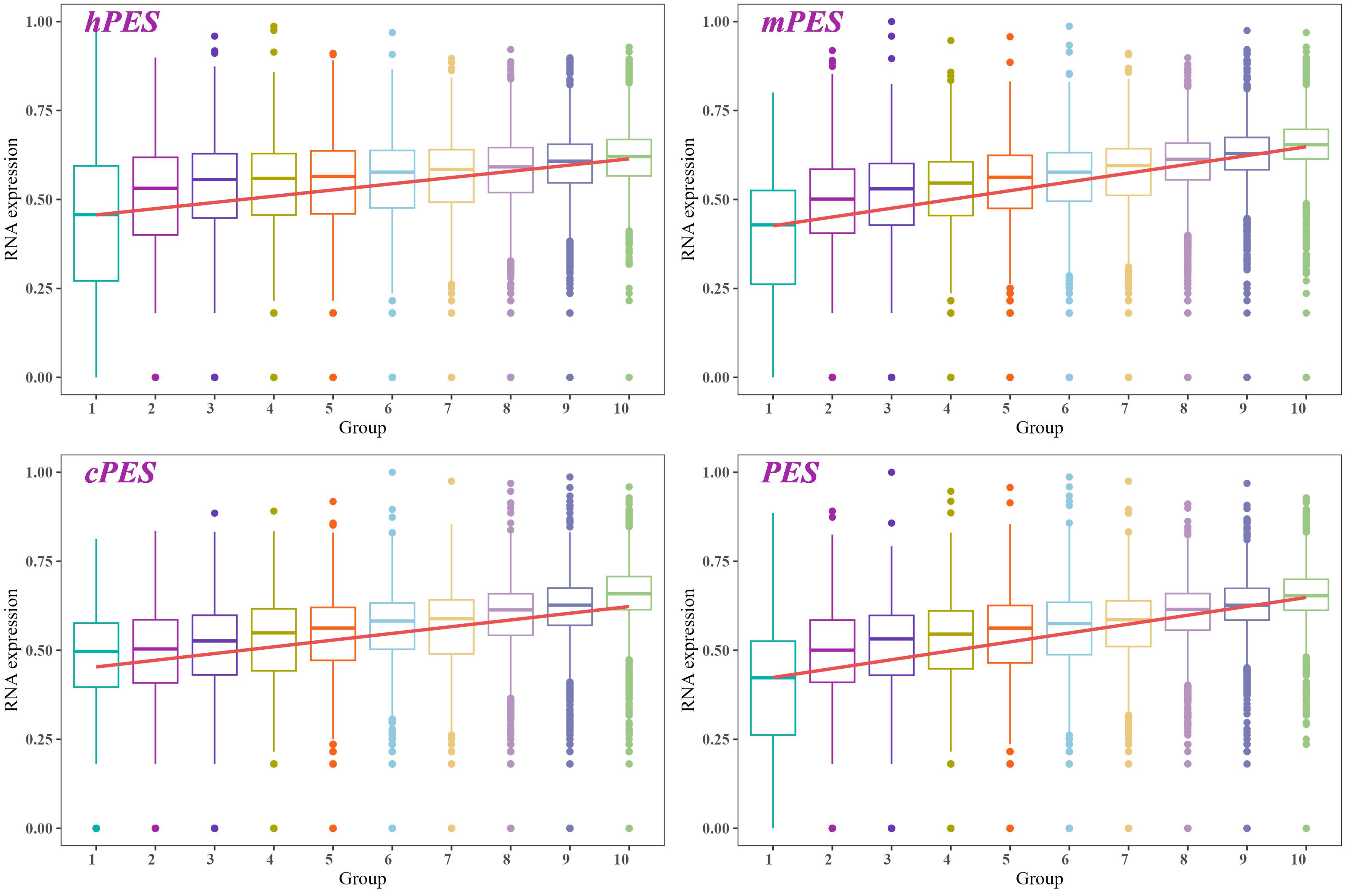

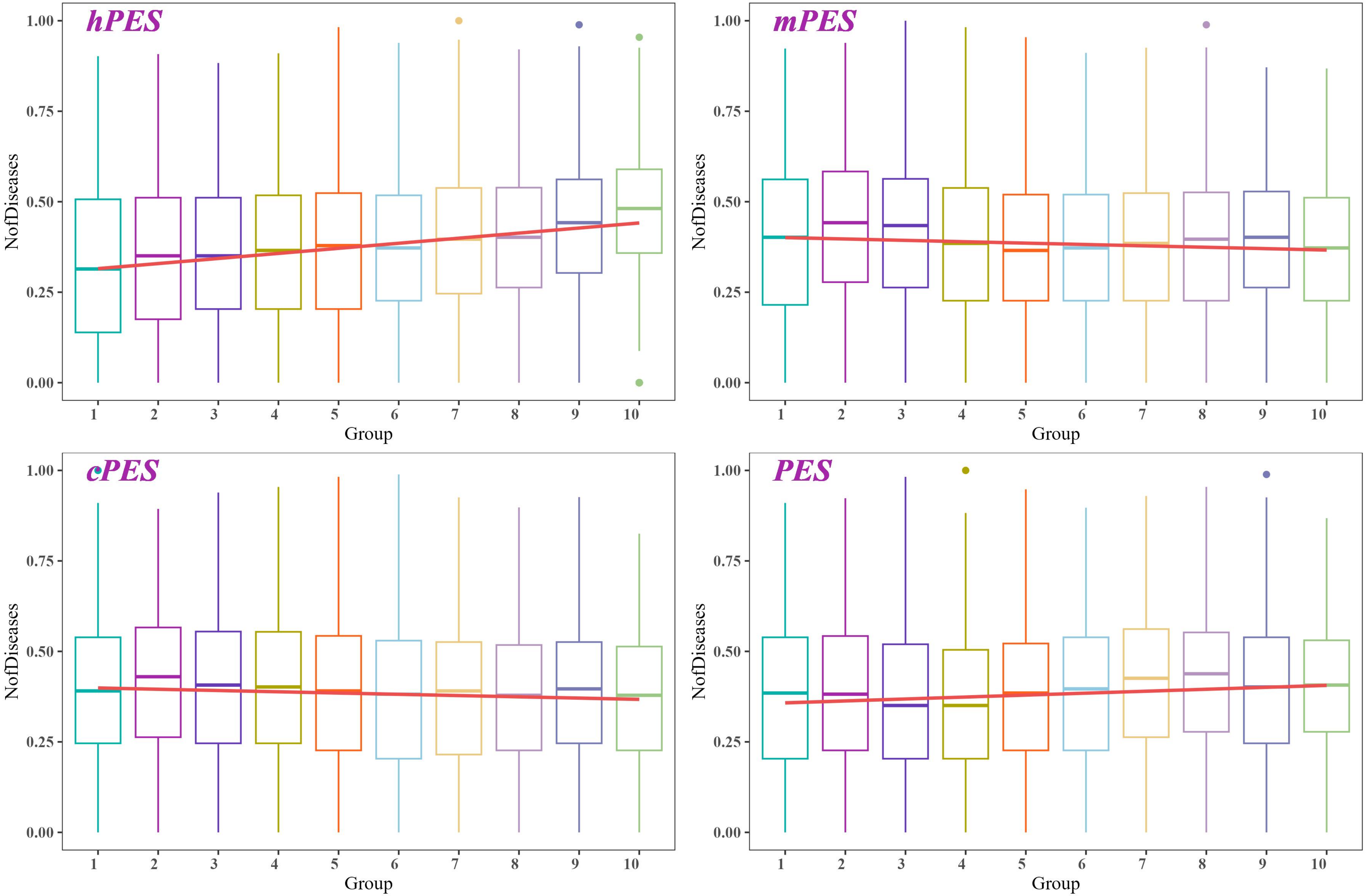

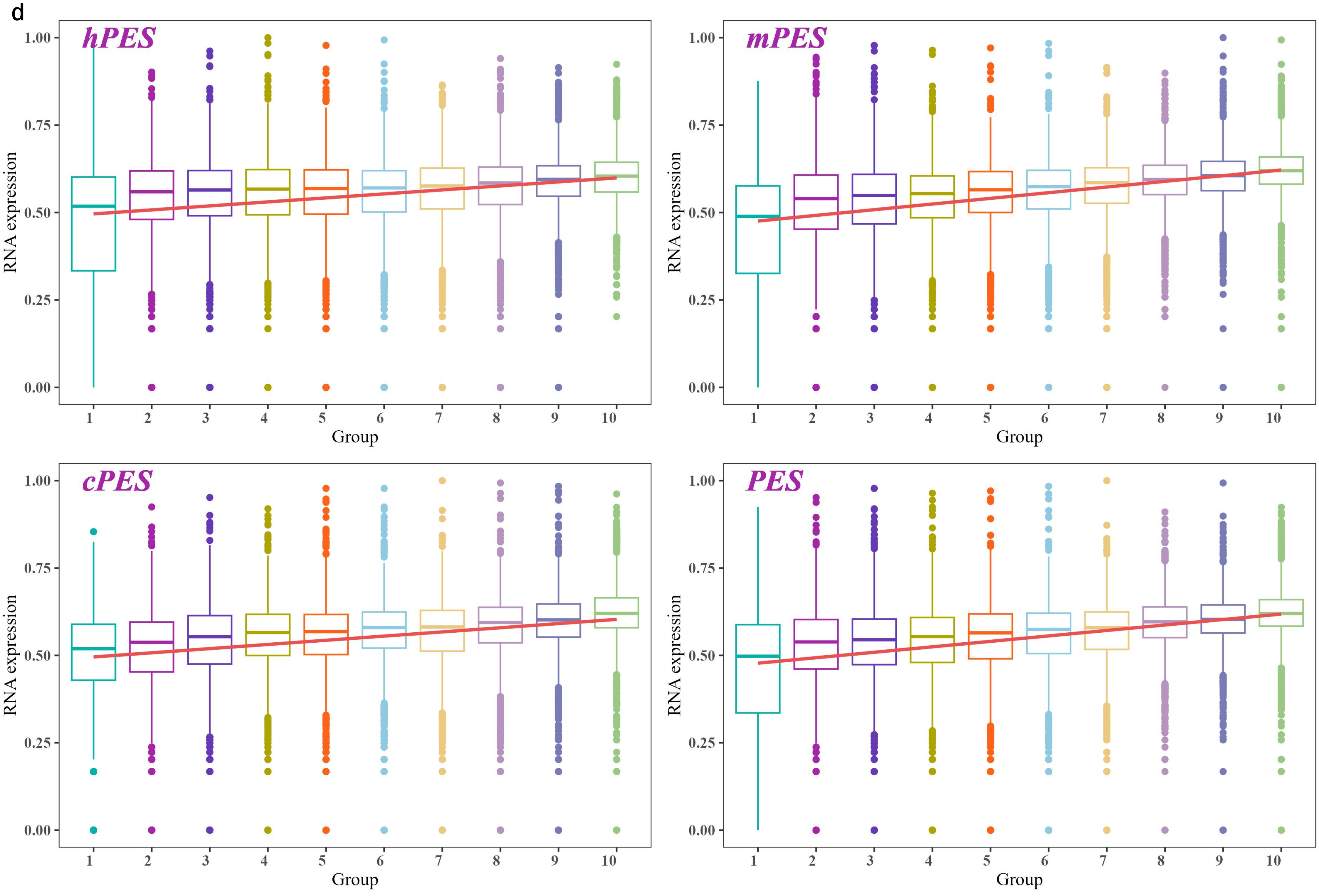

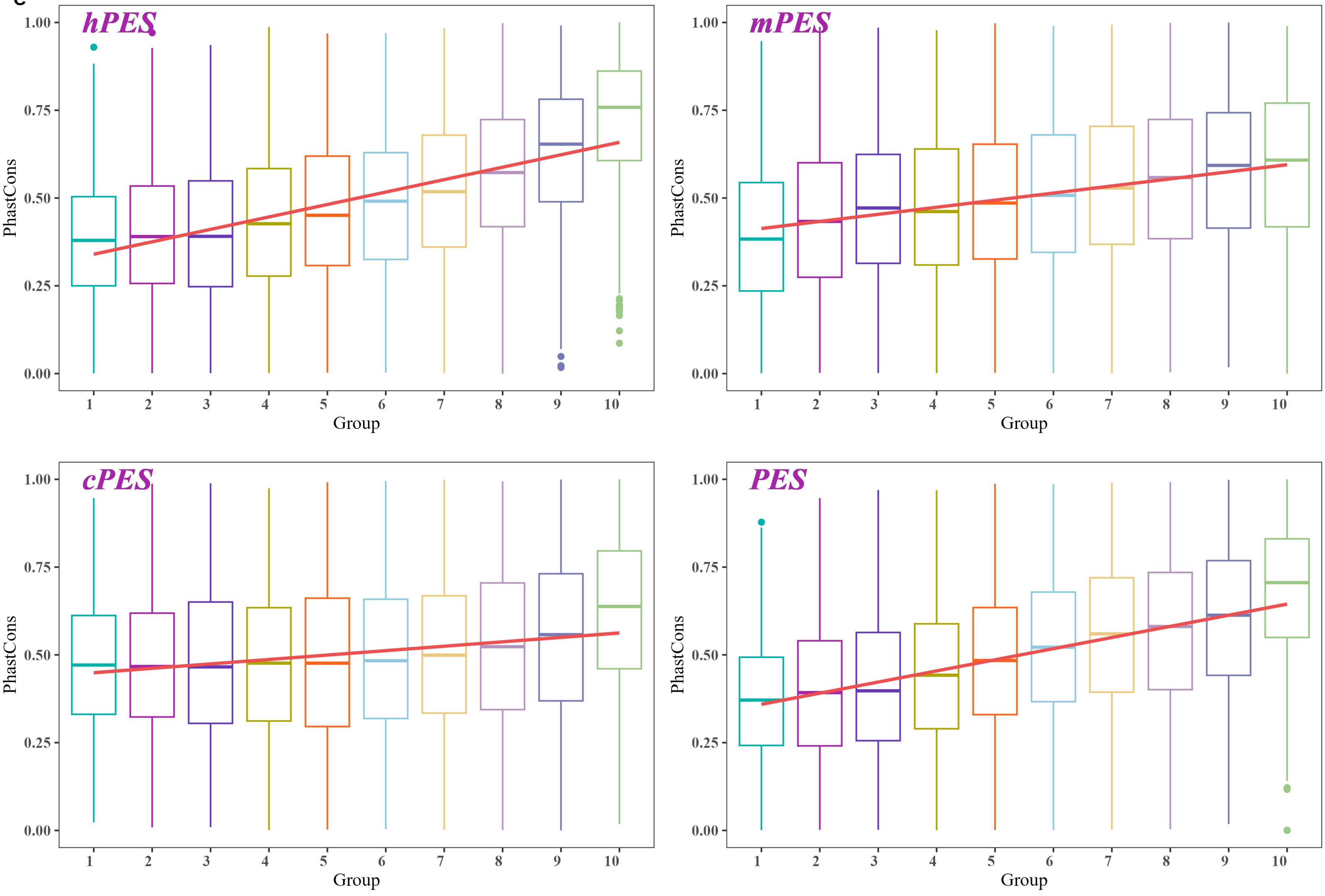

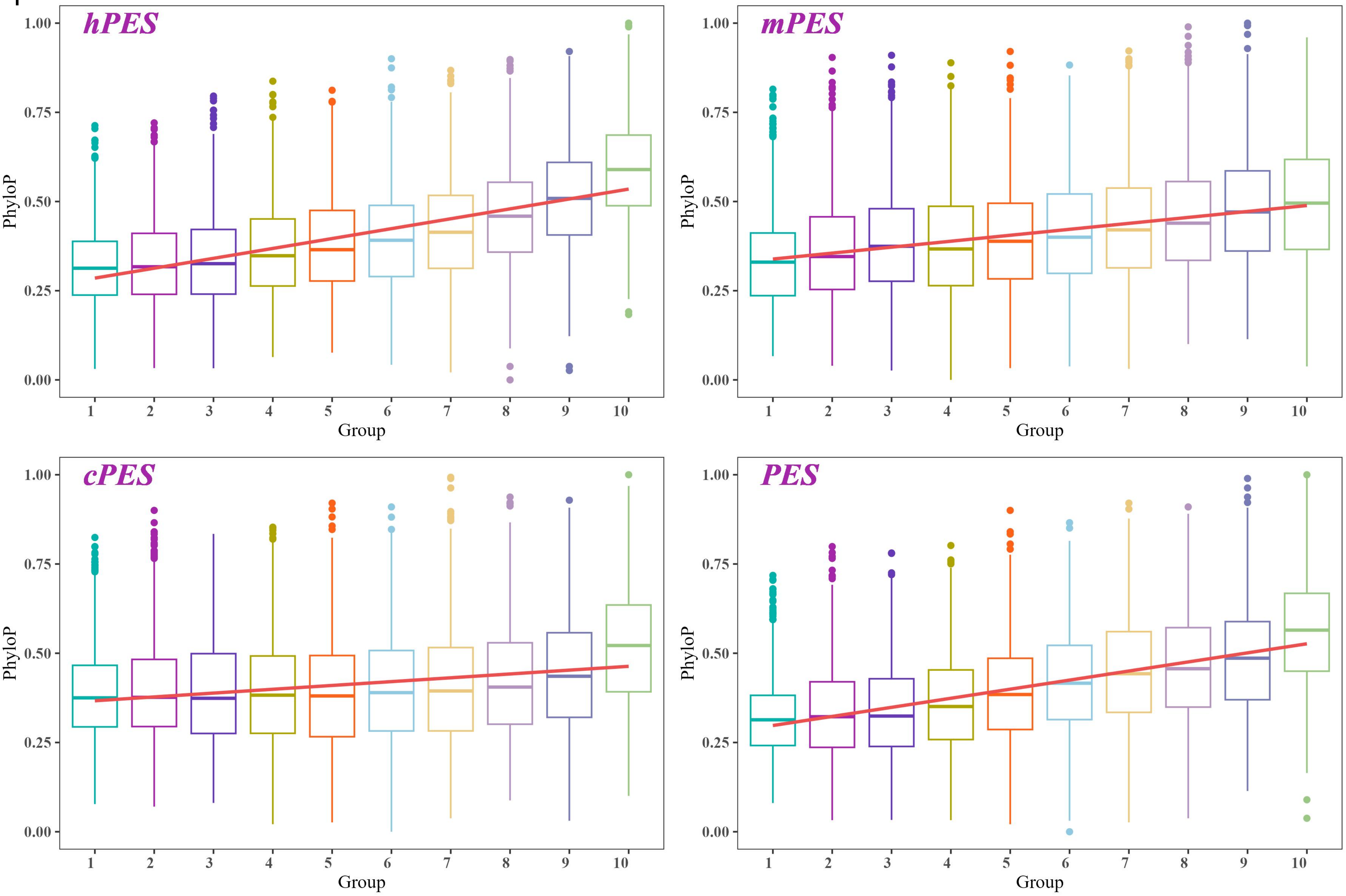

### Supplemental Figure 2

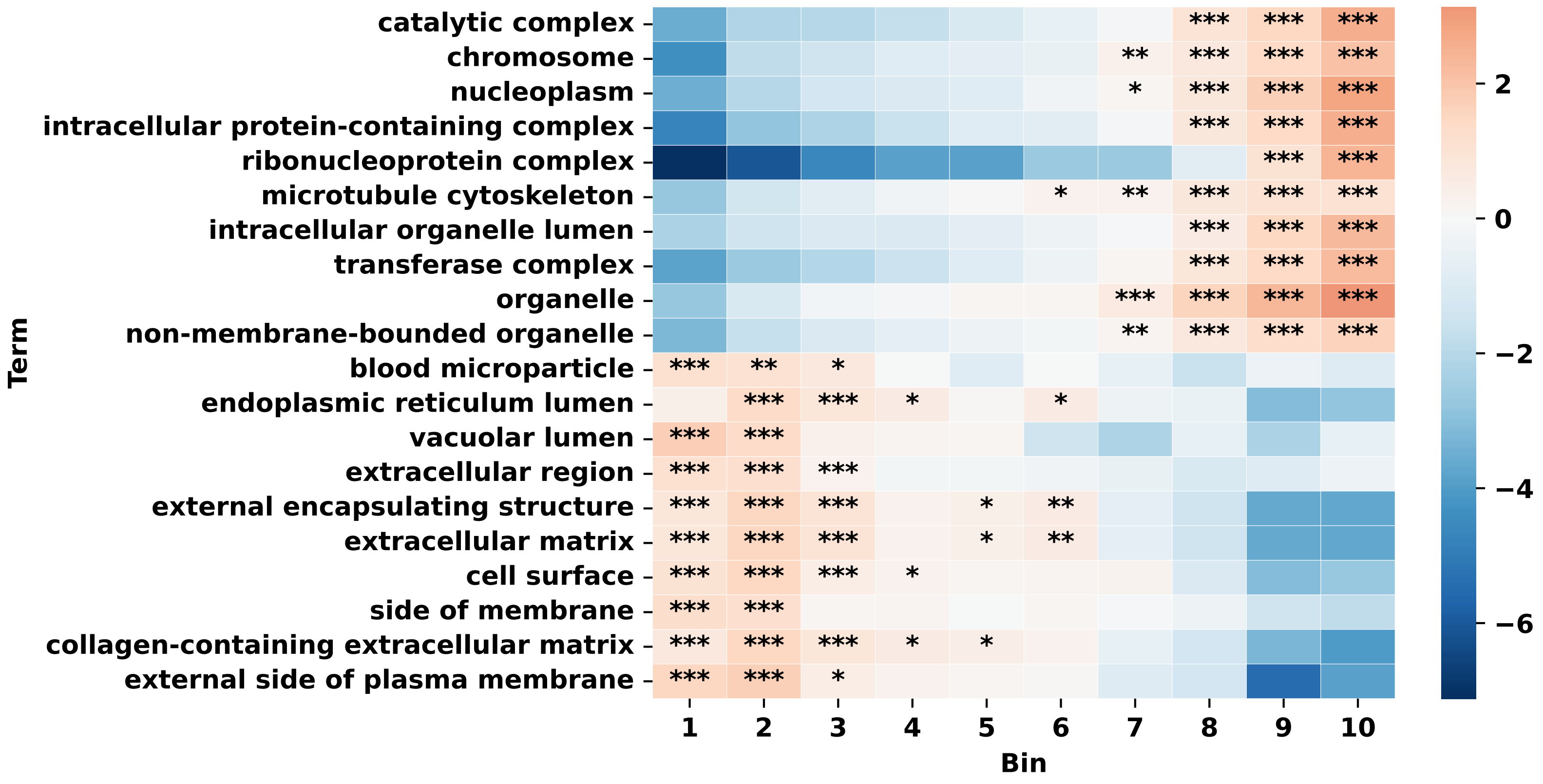

### Supplemental Figure 3

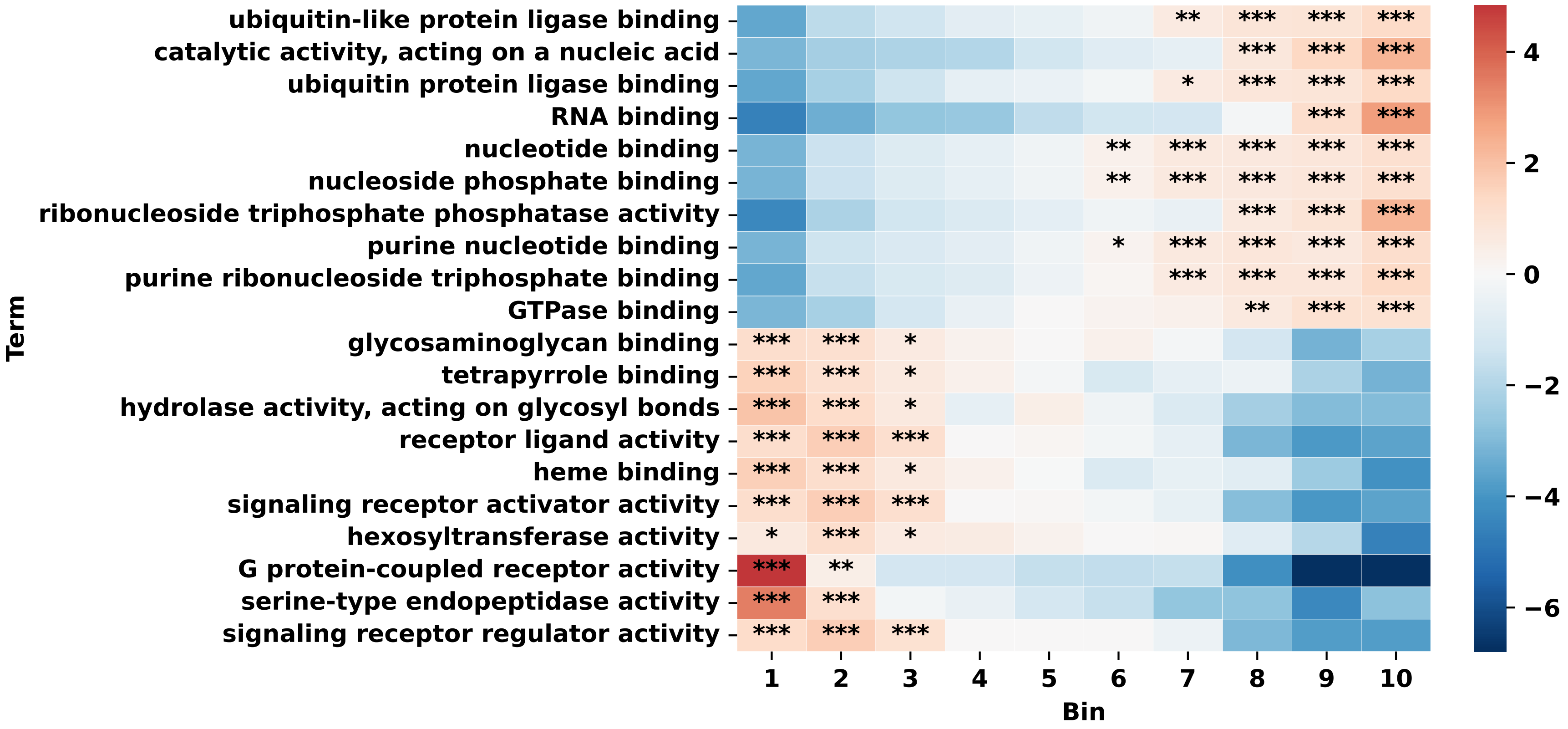
